# Can biocrust moss hide from climate change? Fine-scale habitat sheltering improves summer-stress resistance in *Syntrichia caninervis*

**DOI:** 10.1101/2023.11.06.565694

**Authors:** Theresa A. Clark, Alexander Russell, Joshua L. Greenwood, Dale Devitt, Daniel Stanton, Lloyd R. Stark

## Abstract

**Premise:** Mosses provide many ecosystem functions and are the most vulnerable of biocrust organisms to climate change due to their sensitive water relations stressed by summer aridity. Given their small size, moss stress resistance may be more dependent on fine-scale habitat than macroscale (climate and community), but this sheltering role of habitat (i.e. habitat buffering) has never been compared to macroclimate and may have important implications for predicting critical biocrust refugia in changing climates.

**Methods:** We located three populations of a keystone biocrust moss, *Syntrichia caninervis*, spanning 1200-m of altitude comprising three plant communities (elevation-plant zones) in the Mojave Desert. We stratified 96 microsites along three habitat aridity gradients: elevation-zone, topography (aspect), and microhabitat (shrub proximity). We estimated summer photosynthetic stress (F_v_/F_m_) and aridity exposure (macroclimate, irradiance, and shade).

**Results:** Microsite aridity exposure varied greatly revealing exposed and buffered microhabitats in all elevation-zones. Moss stress did not differ by elevation zone despite the extensive macroclimate gradient, failing to support the high-elevation refugia hypothesis. Instead, stress was lowest on N-facing slopes and microhabitats with higher shrub shading, while the importance of (and interactions between) topography, irradiance, and shade varied by elevation zone.

**Conclusions:** We demonstrate fine-scale habitat buffering is physiologically more protective than high-elevation climate, and thus, may allow some mosses to hide from the brunt of climate change in widespread microrefugia throughout their current ranges. Our findings support a scale-focused vulnerability paradigm: microrefugia may be more important than macrorefugia for bolstering biocrust moss resistance to summer climate stress.

## Introduction

Biocrusts (i.e., biological soil crusts) are diverse networks of cyanobacteria, fungi, algae, lichens, mosses, and liverworts weaving together the upper few centimeters of topsoil in arid and semiarid environments globally (Belnap 2003). Where sufficient moisture or shade exist, mosses are often the larger members of these soil communities and drive many biocrust ecosystem services (Bowker et al. 2013) mediated by their large (typically 1 – 10 cm^2^), absorbent colonies or “patches” that can increase water infiltration and retention (Lafuente et al. 2018), buffer temperature (Xiao et al. 2015), increase soil fertility (Belnap 2003), shelter microbiota (Abed et al. 2019, Fisher et al. 2020), store carbon (Elbert et al. 2012), and prevent erosion (Stovall et al. 2022). However, biocrust mosses are predicted to have more dramatic responses to climate change than most other poikilohydric biocrust species because of their requirements for higher shade and moisture during hydration periods (i.e., “hydroperiods”) when the plants are metabolically active (He et al. 2016, Rodriguez-Caballero et al. 2018, Weber et al. 2018, Ladrón de Guevara & Maestre 2022). Such sensitivity in biocrust mosses has been documented in multiple field and laboratory experiments simulating extreme climate stress (e.g., rapid drying events), which has caused stress, tissue damage, or lethality (e.g. Brinda et al. 2011, Stark et al. 2011, Reed et al. 2012, Zhang et al. 2016, Greenwood et al. 2019, Coe et al. 2020). Moreover, ecophysiological models predict that climatic change may greatly reduce biocrust moss biomass via reduced productivity and increased mortality (Coe and Sparks 2014). Collectively, such impacts on biocrust moss resilience and productivity may reduce their diversity and ecosystem services in changing climates (Ladrón De Guevara & Maestre 2022).

High-elevation habitat such as mountains and plateaus offer cooler temperatures and higher humidity than lower elevations in drylands and thus are predicted to offer oasis to many plants as climate warms and becomes drier (Kelly & Goulden 2008). Such macroclimate gradients along broad altitudinal ranges drive zonation in the dominant vegetation community (elevation zones). High-elevation communities are already documented to support greater moss abundance and diversity in several drylands with extensive aridity-elevation gradients exceeding hundreds of meters (e.g. Nash et al. 1977, Seppelt et al. 2016, Clark 2020). Such elevation zone gradients may, therefore, function as high-elevation refugia for dryland mosses, assuming species have time and sufficient propagules to migrate upward (Zanatta et al. 2020).

An alternative or co-occurring possibility is that mosses could “hide from climate change” in sheltered microrefugia that offer optimal microhabitat within a species’ present range (Ashcroft 2010). The microscale of dryland moss colonies or “patches” are often less than 5 cm^2^ (Nash et al. 1977, Clark 2020) such that fine-scale habitat structure within sites may be equally or more important to moss resiliency than macroscale climate gradients. Fine-scale habitat such as pole-facing slopes and rock surfaces and shade vegetation have been shown to reduce local extremes in temperature, incident radiation, and evaporative demand adjacent to dryland mosses (Alpert 1988, He et al. 2016, Li et al. 2018, Ladrón de Guevara & Maestre 2022). The habitat buffering hypothesis predicts that such sheltered microhabitats will “buffer”, or reduce, climate stress for many organisms (Williams et al. 2008, Scheffers et al. 2014, Shi et al. 2016). Moreover, mosses are small enough to be sheltered by multiple scales of habitat in additive or interactive ways, which we will call *multiscale* habitat buffering. In drylands, there is a prevalence of topographically complex, multiscaled habitat for biocrust; for example a habitat that is shaded by fine-scale vegetation on a north-facing slope within a drainage basin (e.g., Williams et al. 2013, Pietrasiak et al. 2014, Bowker et al. 2016). There is potential that such scales of habitat structure may interact to create strongly sheltered biocrust habitats in future climates.

Predicting the most important scales of habitat buffering to biocrust mosses in changing climates can begin with studying present-day stress and mortality patterns across scales of habitat structure relevant to these mosses. Scales known to influence their distributions include macro-, meso-, and microscale environmental features (Bowker et al. 2016). The mesoscale includes within-site topographical variation on the scale of meters where changes in aspect, slope, and hydrological position can alter soil-water dynamics important to biocrust water availability, such as location relative to a drainage. Microscale variation can include microtopographical shading driven by proximity and azimuth to surrounding vegetation and rocks. Such studies will elucidate the types of refugia, macro-or microrefugia (Ashcroft 2010), that are more probable for small, photosynthetic organisms. Measuring climate stress estimates using gas exchange or chlorophyll fluorescence, will also provide spatially explicit data that can be used to strengthen predictions for future moss abundance distributions in drylands (Coe & Sparks 2014).

We sought to test the high-elevation refugia and habitat buffering hypotheses in the biocrust moss, *Syntrichia caninervis* Mitten, a member of the acrocarpous moss family, Pottiaceae. This keystone biocrust species is understood to be one of the most broadly distributed and ecologically important biocrust mosses in western North America, North Africa, and Asia (e.g., Coe et al. 2012, Seppelt et al. 2016, Ros et al. 1999, Zhang & Zhang 2019). With a broad geographical and altitudinal distribution in the American Southwest (BFNA 2007), *S. caninervis* provides an ideal model species to study summer stress resistance along multiscaled gradients in habitat structure. The Mojave Desert is the most climatically extreme part the species’ North American range where resident populations likely exist at or near physiological thresholds of precipitation minima and temperature maxima (e.g. Stark et al. 2009). Summer presents the highest risk for moss mortality when combinations of extreme desiccation (i.e., cellular water potentials < -400 mPa) interrupted by small rain events have been shown to prevent *S. caninervis* from achieving positive carbon balance during summer hydroperiods (Coe et al. 2012b). Such carbon deficits can lead to full-patch mortality and suggest *S. caninervis* may be increasingly threatened by continued climate change. Current climate trends and predictions in this most arid North American desert include smaller summer rain events and increased drought intensity and variability (Seager et al. 2007, Seager & Vecchi 2010, Zhang et al. 2021).

### Objectives

Our specific objectives in the Desert National Wildlife Refuge (DNWR) along a ∼1200-m elevation zone gradient in Nevada were to: (1) locate 96 *S. caninervis* microsites spanning three nested scales of habitat structure by surveying three elevation zones (macroscale), three topography zones per site (mesoscale), and three microhabitat types per zone (microscale), (2) estimate moss habitat buffering at each microsite (shade time and potential insolation), (3) use in-situ chlorophyll fluorescence to estimate end-of-summer moss photosynthetic stress and mortality, and (4) elucidate the most important scale of habitat structure by testing for stress patterns by habitat scale and modeling the relationship between habitat buffering proxies (elevation, potential insolation, and shade) and summer stress in *S. caninervis*. This natural experiment allowed us to test relationships between three scales of habitat structure and habitat buffering with summer stress resistance in three elevation zone populations of a biocrust moss to gather evidence to either support or reject the high-elevation and habitat buffering hypotheses in one of the harshest environments for which this species occurs globally, the Mojave Desert. Results have important implications for predicting and effectively monitoring the response of this and similar biocrust species to future climate change in drylands by using scale-appropriate and spatially explicit ecological models (Gignac 2001, Dunning et al. 1995).

## MATERIALS AND METHODS

### Elevation zone sites

The Desert National Wildlife Refuge (DNWR) is a large (6,430 km^2^) topographically and biologically diverse basin and range landscape in the eastern Mojave Desert of southern Nevada. DNWR is home to the Sheep Range EPSCoR-NevCAN transect (*Established Program to Stimulate Competitive Research - Nevada Ecohydrological Climate Assessment Network;* Mensing et al. 2013), a set of five climate stations spanning 2000 m of elevation and located in one of the five Mojave elevation zones (**Fig. 1a**), all of which have been floristically characterized without mention of bryophytes (Ackerman 2003, NCCP 2018). Regional soils are limestone-derived, highly calcareous, and range from low-organic to relatively high organic content at high elevations.

**Figure 1.**
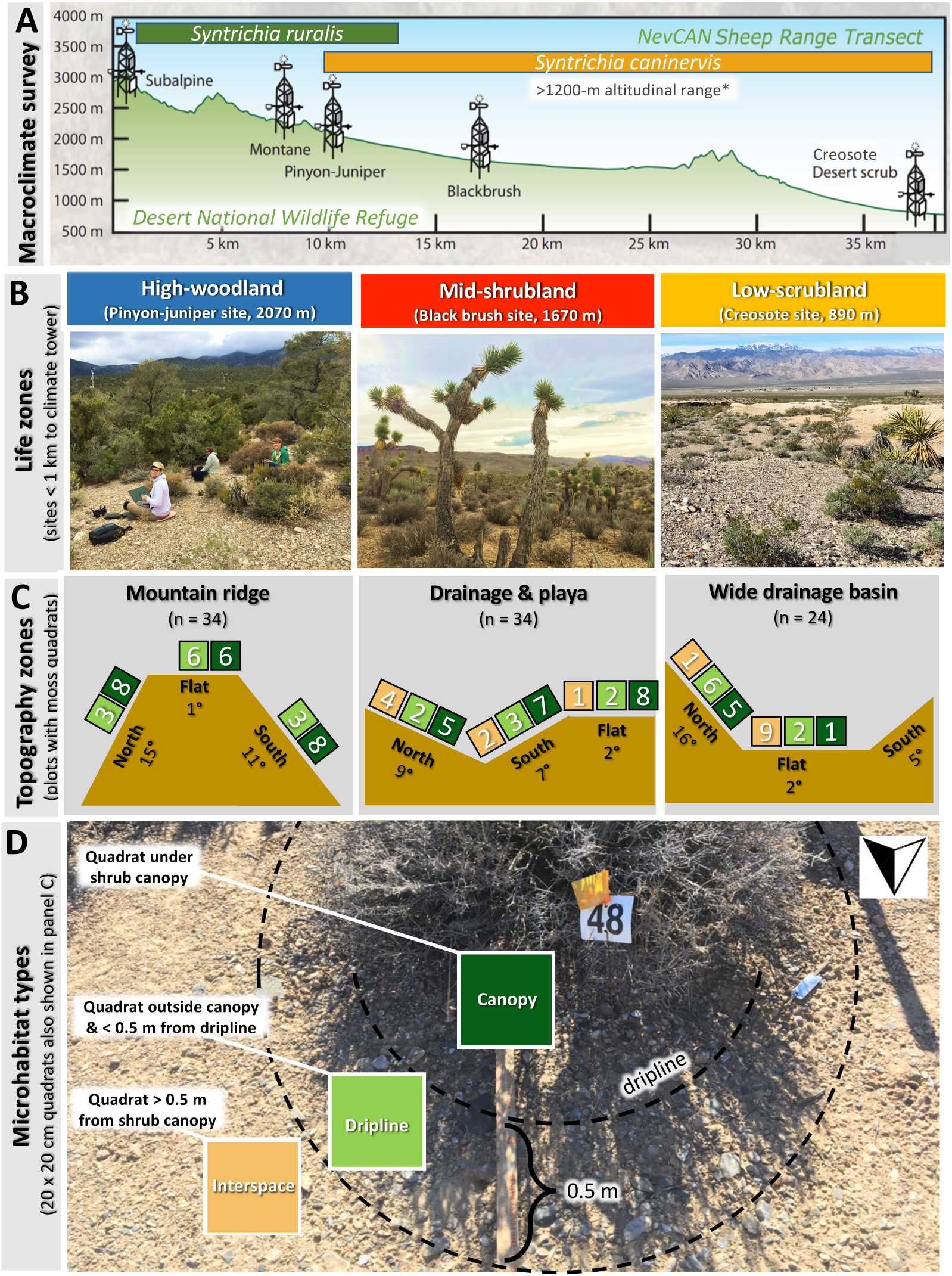
Distribution survey & sampling of populations across nested habitat gradients. **A.** Elevation range surveyed for *S. caninervis* in the Sheep Range of the Mojave Desert National Wildlife Refuge (DNWR). EPSCoR-NevCAN climate towers are shown in each elevation zone (see text). Final sample of 92 microsites (20 x 20 cm quadrats enumerated in **C**) spanned three aridity exposure gradients: elevation zones (**B**), topography zones (northerly, southerly, or flat transects, **C**), and microhabitat shade types (**D**). For each topography zone in **C**, the slope and microhabitat frequency are shown and missing squares indicate *S. caninervis* was not found for a given zone or microhabitat type. See Figure 3 for example quadrats.

The lowest elevation desert scrubland site (890 m, N36.435345, W115.355850; “Low-scrubland”) and surrounding landscape are characterized by an open salt basin interrupted by shallow, calcareous drainages with gentle slopes and occasional steep ravines 1 – 2 m deep. Excluding drainages, the soil is covered almost entirely by desert pavement (e.g. Pietrasiak et al. 2014) with well-spaced shrubs (>2 m apart) and (**Fig. 2a**). The blackbrush-Joshua tree (*Coleogyne ramosissima and Yucca brevifolia*) elevation zone site (1680 m, N36.51723, W115.16191; “Mid-shrubland”) is situated in the center of an intermountain basin divided by drainages that range from ∼1 – 3 m deep. The ground is nearly covered by desert pavement, moderately spaced shrubs (<2 m apart), and widely spaced succulents (**Fig. 2a**). The pinyon-juniper woodland site (2065 m, N36.572808, W115.204060; “High-woodland”) is at the base of the Sheep Mountains on one of the deeply divided ridges with steep, rocky slopes (>3 m tall, ∼10° - 15°). The soil is nearly covered by loose gravel with a dense community of short and tall shrubs and the dominant well-spaced pygmy conifers (*Pinus monophyla* and *Juniperus osteosperma*; **Fig. 2a**). The Montane site (2320 m, N36.590255, W115.214166) has an open-canopy of *Pinus ponderosa* and well-spaced shrubs (>3m apart). The highest-elevation Subalpine site (3015 m, N36.657641 W115.200777) in the NevCAN transect is a nearly closed-canopy mixed-conifer forest (*Abies* concolor, *A. lasiocarpa*, *Picea englemannii*, and *Pinus longaeva*) with calcareous, rocky, organic soils (Ackerman 2003).

**Figure 2.**
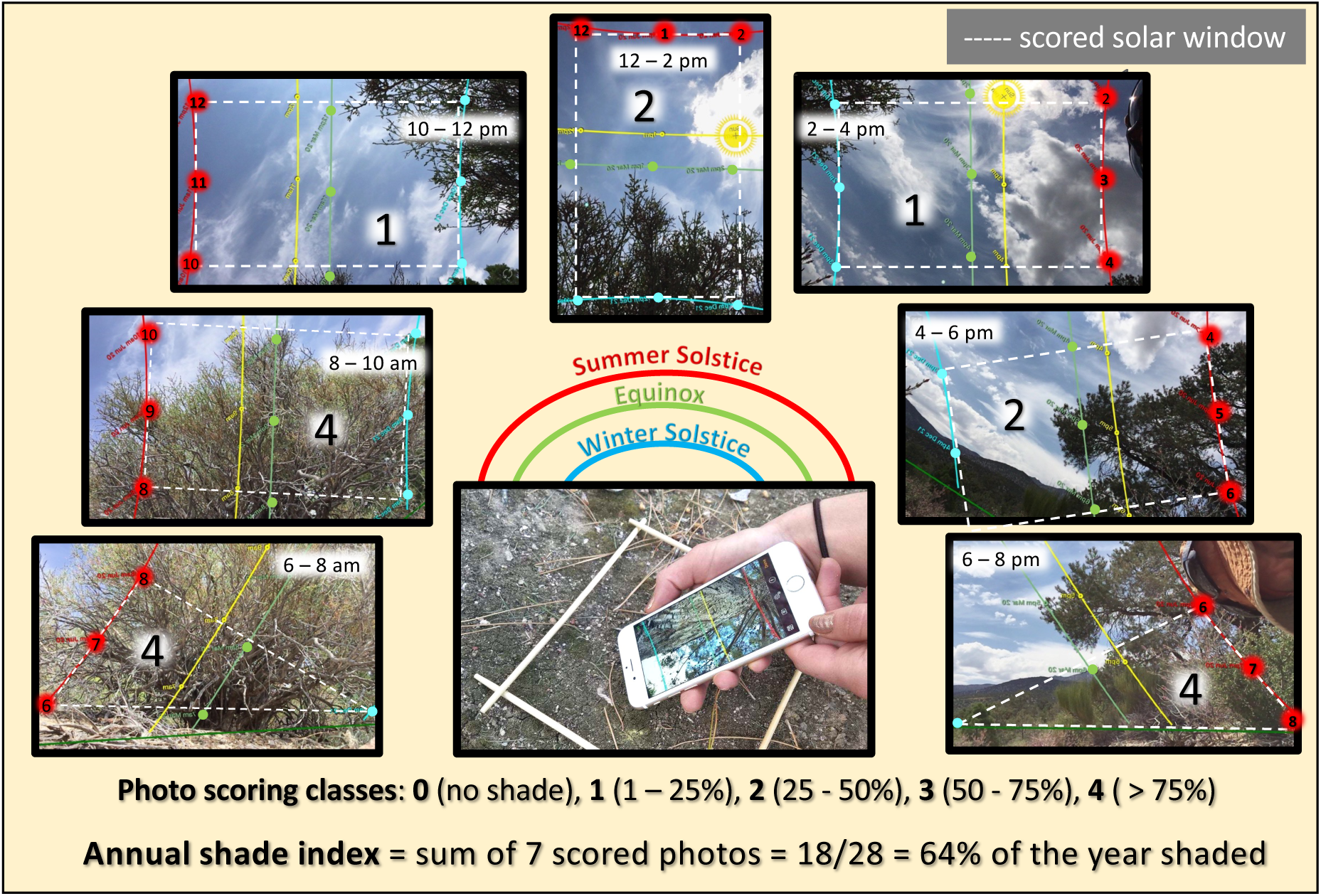
Sun Seeker© shade time method. Microsite annual shade time was quantified using seven photos taken on a smartphone using the *Sun Seeker*© *Solar AR Tracker* app. The phone was held approximately 1 cm above the center of each microsite quadrat (***center photo***) and the photos encompass the solar arc from 6 am to 8 pm (***red circles***), which collectively captures the annual solar window from summer solstice (***red lines***) to winter solstice (blue lines) for a single microsite. Photos are scored using a 5-scale percent shade class by assessing the area of all shade objects intersecting each 2-h solar window (***white dashed boxes and triangles***). Percent annual shade time is the sum of classes for all seven photos divided by 28, the maximum possible (i.e. for a habitat shaded 75-100% of the year). The microsite shown here is shaded 64% of the year ((4 + 4 + 1 + 2 + 1 + 2 + 4 = 18)/28 = 0.64 x 100). Note: the center photo in this figure showing the smartphone position is not located at the microsite of the illustrated solar window.

### Climate metrics

To compare mean annual and summer climate where we located *S. caninervis* along the NevCAN-DNWR elevation zone transect, we acquired NevCAN daily means for temperature, humidity, irradiance, and precipitation from 2011 – 2018 (DRI, 2020). We calculated the 7-yr mean annual air and soil temperature, percent relative humidity (RH), wind speed, and soil moisture. We calculated summer 2017 climate means to include Mojave hot-season months preceding our moss tissue collection, which took place at the beginning of November (6/1/17 – 11/6/17; **Table 1a**).

**Table 1.**
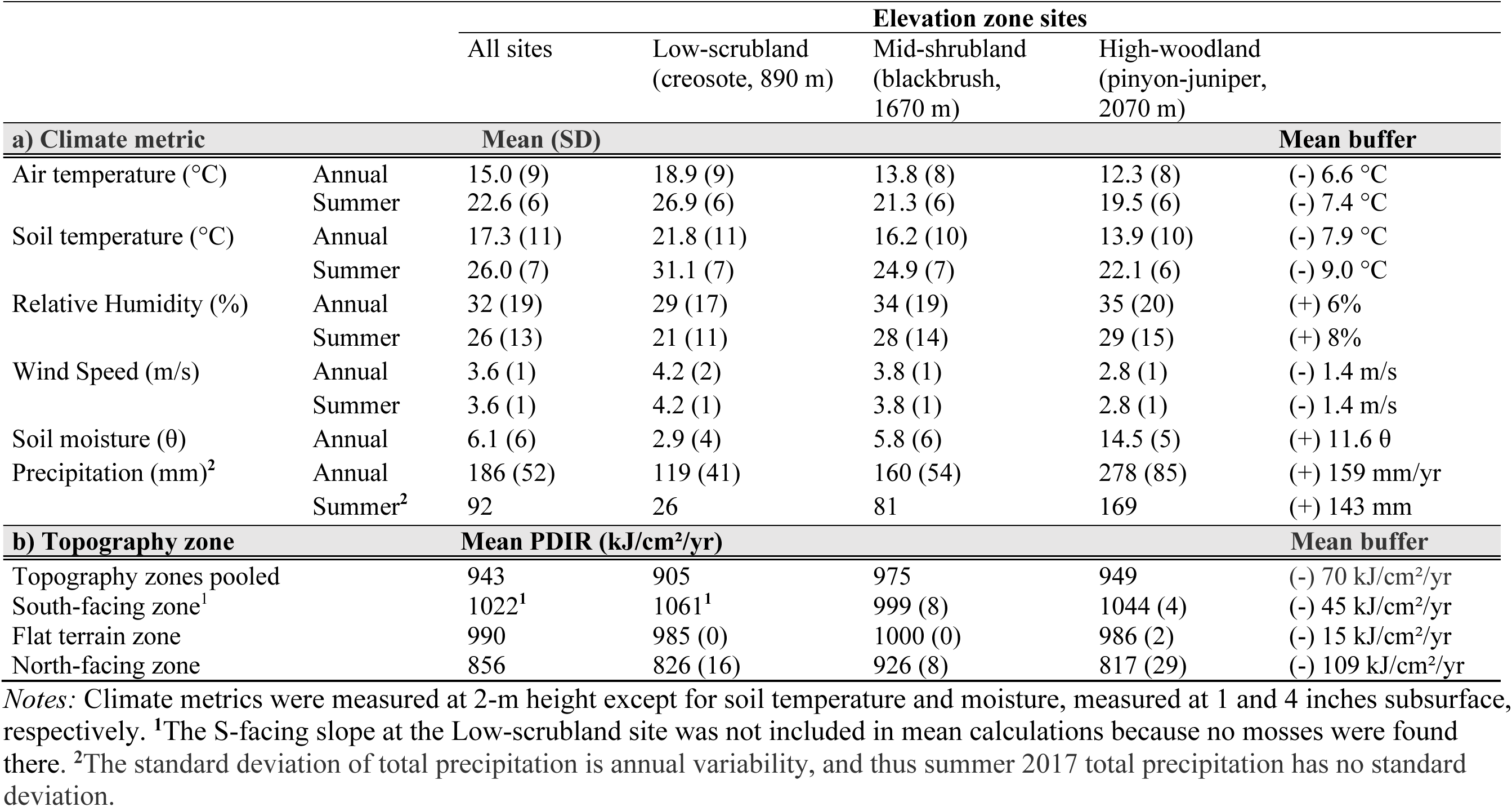
Habitat buffering on three scales (elevation zones, topography zones, & microhabitats) Macroclimate measured by the NevCAN station near each elevation zone site supporting *S. caninervis* in DNWR. **(a)** Annual (2011 – 2018) and Summer (6/1/17 – 11/6/17) daily climate means ± (SD). The maximum elevation zone buffer (**Mean buffer**) is the maximum deviance between the three life-zone means for the respective period. **(b)** Mean ± (SD) potential direct incident radiation (***Mean PDIR***) for elevation zone sites and topography zones (see **Fig. 1**). The greatest topography zone buffer (**Mean buffer**) is the greatest mean deviance between all pairwise comparisons between topography zones for each row. Topography zones varied in their sample sizes (see Fig. 1c); SD = 0 occurred on two topography zones having nearly identical exposure across microsites.

### Aridity exposure survey: elevation zones, topography zones, & microhabitats

To determine the macroclimate exposure of *S. caninervis* in DNWR across the three habitat gradients, we surveyed for species occurrence within a 1-km radius of each elevation zone climate tower (**Fig. 1a& b**); when we found the species in a elevation zone, we surveyed three topographical exposure zones (40 x 10 m plots) selecting the most northerly-facing, southerly-facing, and flat terrain nearest the tower. Selected topography zones varied in their hydrological positions, including association with drainages, uplands, or mountain ridges (**Fig. 1c**). Within each topography zone, we surveyed for moss occurrence in three microhabitat shade classes (“*microhabitat types*” hereafter): (1) high-shade *canopy habitats* partially or fully under shrub canopies, (2) low-shade *interspace habitats* ≥ 0.5 m from the outer edge of the shrub canopy (the “canopy dripline”), and (3) intermediate-shade *dripline habitats* located within a 0.5 m marginal band from the shrub dripline; **Fig. 1d**). Any vegetation located to the north (cardinal 315° – 45°) of moss microsites was ignored in habitat type assignment because it did not contribute to moss shade.

#### Moss microsite selection

We systematically selected 12 microsites per topography zone, attempting to find 4 of each microhabitat type per topography zone, but because this was often not possible (see **Fig. 1c**), the resulting proportion of microhabitats at each elevation zone is a coarse measure of habitat frequency (**S1**). We selected only undisturbed microhabitats having the shrub canopy intact (i.e., not dead or broken off). We systematically centered a 20 x 20 cm quadrat over the highest-density patch of *S. caninervis* in each microhabitat (>3 cm^2^ of *S. caninervis* cover), orienting the quadrat with sides parallel to a North-South axis (**Fig. 1d**). Four shoots were sampled from each quadrat for microscopic species verification; however, mid-way in the study, we removed two microsites after finding (via lab culture) two *Syntrichia* species (*caninervis* and *ruralis* sensu lato) were unknowingly intermixed and possibly measured in the stress assay (Clark 2020; **Fig. 1a**).

### Habitat buffering metrics

Sheltered (i.e. “buffered”) microhabitats for dryland mosses are generally thought to be those with higher humidity, lower temperature, and reduced insolation compared to neighboring local habitats (Alpert 1988). Although such habitat buffering can be measured directly with microenvironmental sensors, large replication (n=96) and sensor installation in delicate biocrust systems is highly invasive; therefore, we measured buffering proxies (e.g. slope, aspect, and shade) for the three scales of habitat structure (utilizing only NevCAN climate tower sensors at each of the NevCAN sites) as follows.

#### Macroscale climate buffering

Elevation zone buffering was estimated at the landscape scale by calculating the mean difference in macroclimate relative to the low-elevation Creosote elevation zone where conditions are most stressful for mosses (i.e. most arid and hot). For example, the elevation zone temperature buffer was calculated as the mean annual decrease in daily temperature between the creosote elevation zone and the higher elevation zones.

#### Mesoscale topographical buffering

We measured aspect (compass cardinal direction) and slope (Suunto PM-5 hand-held clinometer) of the 3 x 3 m^2^ area surrounding each microsite for use in calculating potential direct incident radiation (PDIR). Within elevation zones, we calculated mesoscale topographical exposure as the mean of microsite PDIR in each topography zone. PDIR was estimated with a complex formula that incorporates microsite elevation, latitude, slope, and folded aspect (McCune & Keon 2002, McCune 2007). We observed *S. caninervis* ground cover to be greatest on the N side of shrubs in this ecosystem and thus modified the PDIR equation for use with a N-S rather than NE-SW axis as described in McCune (2007). To estimate PDIR buffering across and within elevation zones, we calculated differences in mean mesoscale topographical exposure. For example, the maximum topographical zone buffer was simply the difference in mean PDIR between topography zones with the highest and lowest mean PDIRs. Mean elevation zone PDIR was simply the average of all topography zone PDIR at each elevation zone site.

#### Microscale shade buffering

To precisely measure fine-scale shade time of microsites, we developed a photographic method using the smartphone app, *Sun Seeker© Solar AR (Augmented Reality) Tracker* (Sydney, Australia), which maps onto the camera view the annual solar window from sunset to sunrise for a given location (e.g., moss microsite). Seven photos taken at any time of the year circumscribe all geographical-time-referenced shade objects from the vantage point of the moss (**Fig. 2**). For each photo, the area of shade objects (i.e., vegetation and topography) inside the solar window was scored using a shade class from 0 (no shade) to 4 (75 – 100% shaded; see classes in **Fig. 2**). For each microsite, the resulting seven shade classes were divided by the total possible score of 28 shade points to yield an annual shade time percentage. Our novel microscale shade metric estimates the percentage of the year a microhabitat is shaded by vegetation and/or topography; we used this sensor-free buffering metric to test differences in mean shade at various scales of habitat structure.

### Field sampling & tissue prep

To measure the summer stress signal of *S. caninervis*, we collected shoots when patches were fully desiccated in late fall (November 6 – 13, 2017; *Permit #84555-17-019 U.S. Fish and Wildlife Service National Refuge System Research & Monitoring*), which is still considered the Mojave warm season. This dormant, dry tissue preserved the summer/warm-season stress signal because we collected prior to any fall precipitation during which some recovery could begin. We collected two to three shoots for every 2-cm^2^ patch area per quadrat using fine forceps (**Fig. 3a-b**). Collections were stored dark and dry at 20°C with 20 – 30% relative humidity, conditions which should not exacerbate stress in this species (Guo & Zhao 2018).

**Figure 3.**
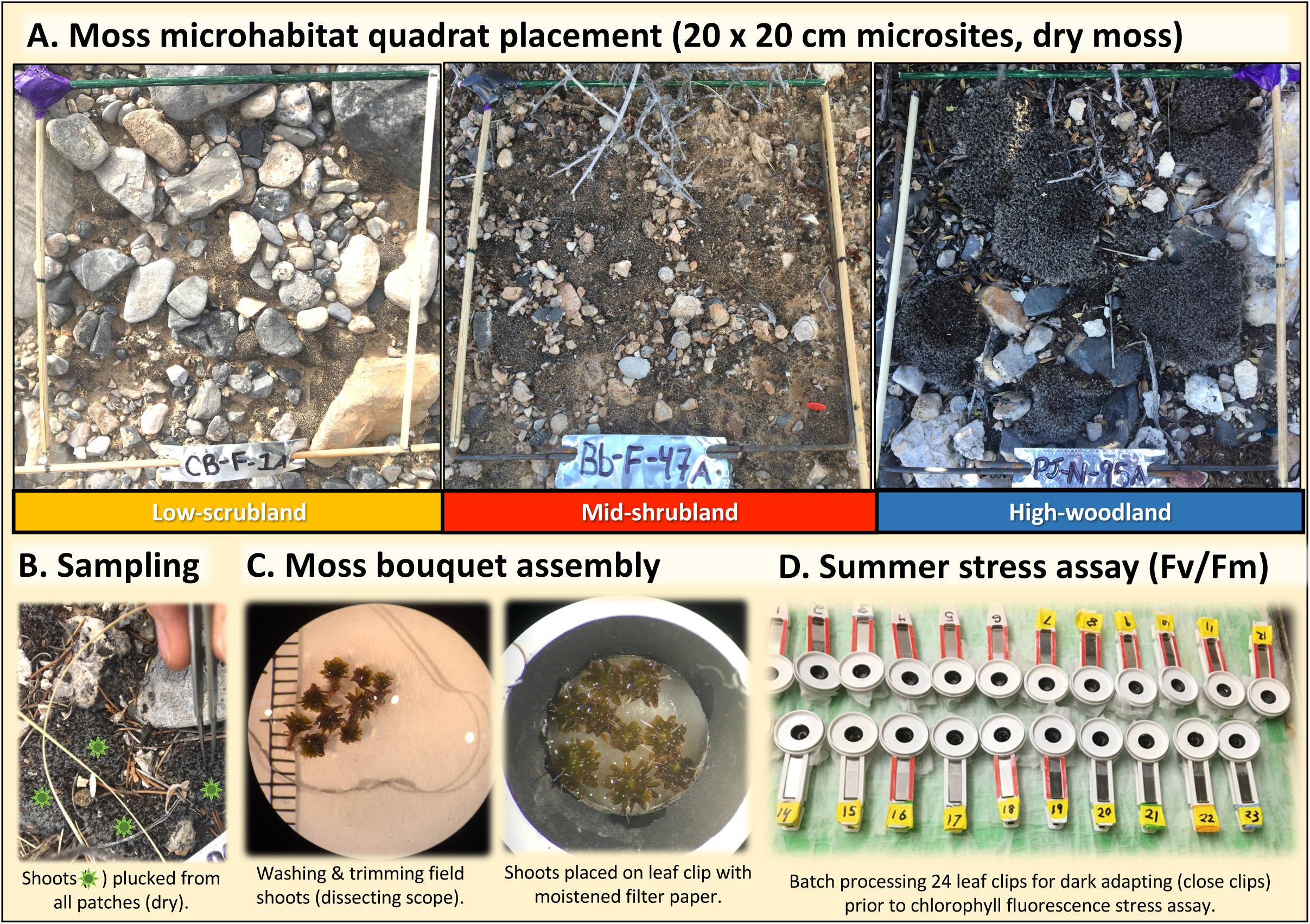
Sampling and stress assay methods for small biocrust moss. Methods for microhabitat selection at three elevation zone sites showing moss abundance gradient (**A**), field shoot sampling (**B**), moss bouquet assembly (**C**), and the photosynthetic efficiency assay (**D**) to test summer stress in a small desert moss.

Before the stress assay, shoots were hydrated with distilled water on a microscope slide and the upper 2 mm were cut and retained to target the living apical “green zone” (Stark 2017) (**Fig. 3b**). Green zones were swirled in a drop of water to remove debris, and placed vertically into a rosette formation on a wetted 7-mm filter paper disc creating a “*moss bouquet”* (**Fig. 3b**). The number of shoots in a moss bouquet (5 – 23) depended on shoot size to maximize microsite representation and standardize leaf area for chlorophyll fluorescence.

### Summer-stress resistance assay (Fv/Fm)

Efficiency of photosystem II (PSII) photosynthesis is commonly measured as a stress proxy in plants via the non-invasive technique, chlorophyll fluorescence (Papadatos et al. 2017), convenient for measurements on small moss species with limited biomass or long-term monitoring (Proctor 2009). We refer to Fv/Fm as a stress proxy because in desiccation tolerant mosses, Fv/Fm responds sensitively and negatively in a near linear fashion to the level of desiccation and light stress induced on species in carefully controlled experiments (e.g. Proctor et al. 2007, Proctor 2001). In almost all cases, the lower the Fv/Fm when such a moss is rehydrated (30 min – 1 hr after desiccation), the more stressful were the conditions during the previous hydration/desiccation cycles (e.g. Coe et al. 2020). This reduced Fv/Fm can be viewed as a photoprotective acclimation to light and low water potentials both of which stress the electron transport chain and induce a photoprotective response (Papadatos et al. 2017). Long-term acclimation to high light and desiccation is the most parsimonious explanation for reduced potential photosynthetic efficiency (Fv/Fm) in desiccation tolerant mosses (when measured on fully hydrated material) regardless of the photoprotective response, which usually is assumed to involve the D1 repair cycle and slowly reversible forms of nonphotochemical quenching (Proctor et al. 2007). Therefore, our photosynthetic stress proxy, Fv/Fm, will be called “stress” in figures and the Discussion.

Our Fv/Fm measurements were performed with a Hansatech FMS-2 modulated fluorometer (Norfolk, England) and modified leaf clips hollowed to accommodate taller shoots (**Fig. 3b & c**). Hydrated moss bouquets were each placed on a folded chemical-wipe and carefully positioned into leaf clips, then placed in a tray of distilled water to maintain full-turgor throughout the 20-min dark acclimation (a period needed to close all active PSII reaction centers). Including 10 min to clean and prep bouquets, we measured chlorophyll fluorescence at 30-min post-rehydration, a standard photosynthetic reactivation timepoint for assessing the photosynthetic stress or vitality of mosses prior to significant recovery from the preceding desiccation event (e.g., Munzi et al. 2019, Hamerlynck et al. 2000, Ekwealor et al. 2021).

A Hansatech script automated our measurement of the dark-acclimated metrics, basal (Fo) and maximum (Fm) fluorescence, during a 0.8-s saturation pulse of 3000 µmol/m^2^/s. Using variable fluorescence (Fv = Fm – Fo), Fv/Fm was then derived, which estimates potential maximum quantum efficiency of photosystem II (PSII) photochemistry of dark-acclimated plants (Baker 2008). Fv/Fm provides a universal physiological indicator of climate stress in plants that is more sensitive and integrative than chlorophyll content (Murchie & Lawson 2013) and requires less tissue and time than gas exchange analysis.

### Analysis

Analyses were performed in *R* (Version 1.447 and 4.2.3; R-Core-Development-Team 2023) using the *tidyverse* package (Wickham et al. 2019) and statistical packages denoted as (*package::function*). We used an exploratory approach with hypothesis testing in which we consider the ecological significance of statistical patterns when *P* is < 0.05 and opt out of arbitrary family corrections for testing our small set of environmental factors (Gotelli and Ellison 2013).

#### Shade buffering & moss stress vs multiscale habitat structure

Although our design involved three categorical habitat predictors of moss stress and shade buffering (elevation zone, topography zone, and microhabitat type), using three-way ANOVA’s to test patterns in mean shade buffering and moss stress were not possible due to missing factor levels (i.e. no S-facing zone in the Low-scrubland and no interspace microhabitats in several topography zones (**Fig. 1c**, **S1**). Alternatively, we conducted six univariate ANOVA’s: three stress tests and three shade buffering tests, one for each habitat factor. We calculated the effect sizes, Eta-squared (equivalent to R^2^ in one-way ANOVA) and Cohen’s *f* (Lakens 2013). When heteroskedasticity was present (determined with the Fligner-Killeen test; Fligner & Killeen 1976), we opted out of transformations, which are known to yield nonsensical predictions of proportion data and hinder interpretability (Warton and Hui 2011). Instead, we used robust Welch’s denominator degrees of freedom corrections appropriate for the approximate normality, unbalanced design, and heteroskedasticity present (*stats::oneway.test*; Welch 1951). Post-hoc tests for Welch’s ANOVA’s were nonparametric Games-Howell multiple comparisons (*rstatix:: games_howell_test*; Kassambara 2023, Ruxton & Beauchamp 2008).

#### Habitat buffering proxies as predictors of moss stress

To explore the relationship between the three scales of habitat buffering (elevation, PDIR, and shade) and summer moss stress, we fit a linear multiple regression model for Fv/Fm (**S2**). Despite small variance inflation factors of 2.07, 1.17, 2.19, for the three predictors, respectively, our model testing procedure revealed instability in parameter estimates and p-values when elevation was included in the model (and no interaction terms were included). Therefore, the final reduced model included only PDIR, shade, and their interaction, centering the explanatory variables to remove structural multicollinearity. Diagnostic residual plots revealed adequate fit (**S3**); however, we acknowledge that beta regression is optimal for modeling such a two-category ratio (Douma & Weedon 2018) and should be used for prediction-focused studies.

#### Elevation zone models

With elevation zone management in mind, we performed a set of hypothesis tests focused on each elevation zone site to explore whether our spatially efficient (i.e. nested) sampling design (**Fig. 1**) of meso-scale topography zones (within-site plots) and micro-scale shade measurements (within-plot microsite quadrats) could explain significant patterns in moss stress. (Note: although ideal, testing these relationships in a single ANOVA model including all elevation zones was not possible due to the multicollinearity between elevation zone (elevation) and shade (as discussed in the habitat buffering model above).

Alternatively, we ran three separate ANOVAs (i.e. ANCOVAs) to test additive and interactive relationships between topography zone and percent shade time (centered before analysis) with moss stress for each elevation zone. ANOVAs were made robust to (a) the unbalanced design (**Fig. 1c**) using Type II sums of squares (Langsrud 2003, Logan 2010) and to (b) unequal variance using a Huber-white heteroskedasticity-corrected covariance matrix (HCCM; White 1980; Long & Ervin 2000) for each model (*car::Anova,* Fox & Weisberg 2019). Post-hoc pairwise comparisons for regression slopes (by topography zone) were not conducted (when the interaction term was significant) because we believe sample sizes greater than 11 – 12 quadrats per topography zone should be collected to produce more accurate fine-scale relationships. The OLS coefficient of determination (*R^2^*) is shown for each final model as an effect size metric; the overall F-test for each HCCM-corrected model was derived from a Wald test comparing the intercept model to the respective HCCM model including only significant terms (*lmtest::waldtest*; Zeileis & Hothorn 2002).

## RESULTS

Results are numbered by objective and statistics are printed in figures if not shown here. We refer to our three scales of aridity gradient sampling (i.e., elevation zones, topography zones, and microhabitat types, **Fig. 1**) as the three scales of habitat structure or habitat buffering, depending on context.

### Aridity exposure across elevation zones (Objective 1)

After extensive surveying within a 1-km radius of each NevCAN climate tower, we located *S. caninervis* biocrust in the lower three sites of the DNWR eco-hydrological gradient spanning 1,180 m of elevation from 890 to 2070 m, while a closely related species, *Syntrichia ruralis* sensu lato, was found primarily at higher elevations (**Fig. 1a**); we will refer to these sampling sites as the Low-scrubland, Mid-shrubland, and High-woodland, respectively (**Fig. 1b**). We located *S. caninervis* on North-facing, South-facing, and flat topography zones at the Mid-shrubland and High-woodland, but it was absent from southerly facing slopes at the Low-scrubland (also absent from upland flats) and was only found on shallow drainage flats and northerly-facing slopes (**Fig. 1c**). In the Low and Mid-elevation sites, *S. caninervis* occurred in all three microhabitat types, but in the High-woodland, shrub interspace habitat did not support high-density *S. caninervis*. Relative habitat frequency changed with elevation: shrub canopy habitat increased, interspace habitat decreased, and shrub-dripline frequency remained similar from low to high elevation (**S1**, **Fig. 1c**). Canopy microhabitats were the most frequent habitat type supporting high-density *S. caninervis* biocrust across elevation zones (54/94 microsites, **S1**).

### Habitat buffering across three scales of habitat structure (Objective 2)

#### Macroscale climate buffering

Mean daily macroclimate differed substantially by elevation zone (**Table 1a**) with higher elevations having cooler temperatures, higher humidity and precipitation, and slower wind speeds. Relative to the Low-scrubland, the High-woodland site was climatically buffered in four metrics: (1) 6.5 times more precipitation, (2) reduced mean daily air and soil temperatures by -7.4°C and -9.0°C, respectively, and (3) increased mean relative humidity by +8% (**Table 1a**).

#### Mesoscale topographical buffering

The nine topography zones collectively represent the mesoscale exposure gradient for *S. caninervis*-dominant biocrust at each elevation zone site and varied in their hydrological position by elevation zone: the Low-scrubland site was in an ephemeral drainage, the Mid-shrubland included a drainage and upland flat, and the High-woodland traversed a mountain ridge (**Fig. 1c**). Of the nine topography zones, there was a 22% reduction from the highest mean PDIR of the S-facing slope of the High-woodland (1044 kJ/cm²/yr) to the lowest mean PDIR of the N-facing slope of the High-woodland (817 kJ/cm²/yr), which created a maximum mean topography zone buffer of 227 kJ/cm²/yr (**Table 1b**, **Fig. 1c**).

Notably, the Low-scrubland had the lowest average PDIR (of 24 microsites) due to its relatively steep 16° N-facing slope.

#### Microscale shade buffering

Percent annual shade time (“shade” hereafter) across the 94 microsites ranged from 21 – 96% and averaged 64 ± 2% (SE) such that most microsites were shaded over half of the year. Elevation zone significantly explained 48% of shade variation (^Welch^F_2, 52_ = 40.6, P < 0.0001); mean shade in the High-woodland was 1.9x greater than in the Low-scrubland (**Fig. 4a**). Mean shade differed little by topography zone when pooling microsites across elevation zones (^Welch^F_2, 58_ = 2.7, P = 0.074, *R^2^* = 0.05, **Fig. 4b**), while microhabitat type explained 67% of variation in shade, which increased 2.5-fold from interspace to canopy habitats, on average (*^Welch^F_2, 44_ = 44.0*, *P* < 0.0001, **Fig. 4c**).

**Figure 4.**
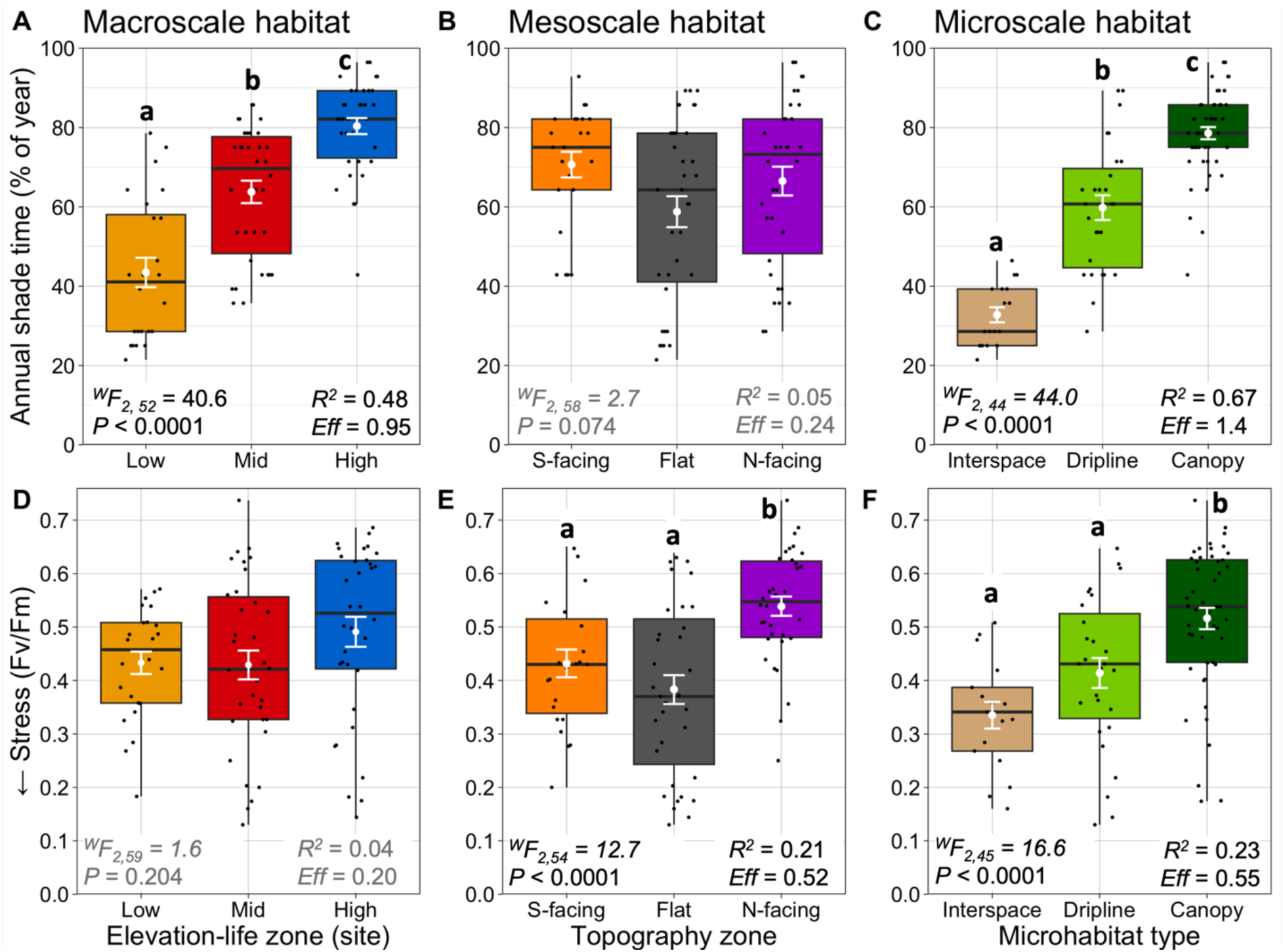
Shade buffering & moss stress across three scales of habitat structure. Boxplots of shade time (top row) and summer stress (bottom row) measured in 92 *Syntrichia caninervis* microhabitats pooled by scale of habitat structure: elevation zone (**A & D**), topography zone (**B & E**), and microhabitat type (**C & F**). Overlays include raw data (black points) and respective means (white points) with SE bars. Welch’s ANOVA results, *R^2^*, and Cohen’s *F* (*Eff*) are shown for each panel. Letters indicate familywise Games-Howell significant differences between groups (*P* < 0.05) in post-hoc testing. Denominator degrees of freedom in F-statistics do not reflect sample size due to Welch’s heterogeneity correction.

### Summer Stress Proxy (Fv/Fm) by elevation zone (Objective 3)

Nearly all *S. caninervis* patches showed visible signs of stress in their darkened, sun-pigmented leaves (**Fig. 5**). The sample distribution of Fv/Fm was left-skewed with a mean of 0.455 ± 0.148 (SE) and ranged from 0.130 at the Mid-shrubland to 0.737 at the Mid-shrubland (**Fig. 6**). Of the 94 mosses, ∼23% were near normal levels (Fv/Fm >0.6) leaving 77% stressed (see *Discussion*). Dividing the typical physiological range of Fv/Fm (0 – 0.85) into four stress categories, we report 7% severely stressed, 25% moderately stressed, and 42% mildly stressed, and 26% unstressed (**Fig. 6**). The seven samples with Fv/Fm < 0.2 (in the lower 25th percentile; **Fig. 6**) were from Flat-zones at all three elevation zones. These outliers included two interspace, three dripline, and one canopy microhabitat and had moderate annual shading ranging from 43 – 68%, except for one interspace microsite at the Low-scrubland that was shaded only 28% of the year. Signs of extreme stress in these samples were evident in chlorotic leaf tips (**Fig. 5b-c**).

**Figure 5.**
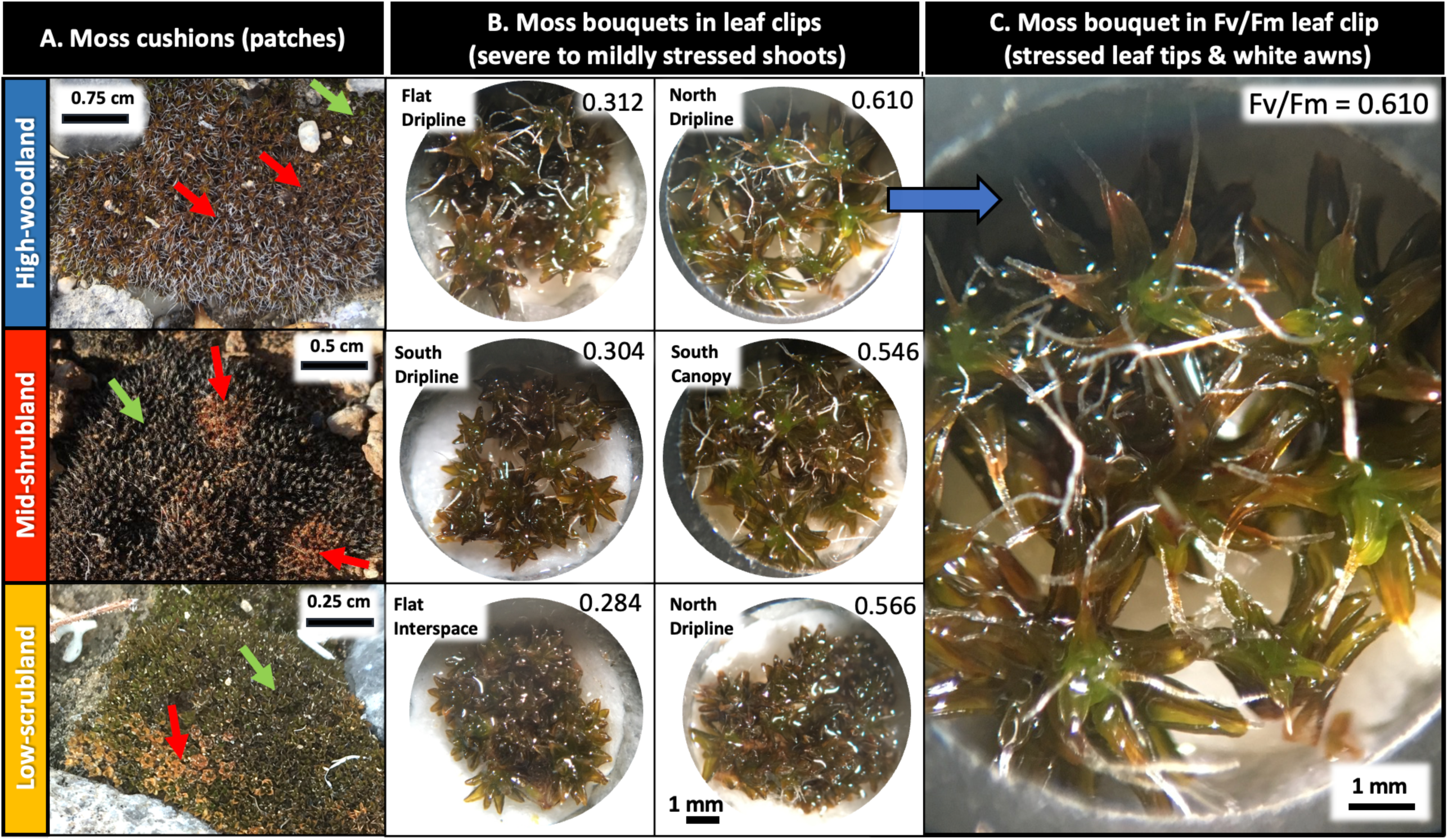
Patch appearance & shoot mortality: healthy & stressed *S. caninervis* in three Mojave Desert elevation zones. *Syntrichia caninervis* patches (i.e. shoot colonies) by elevation zone site (***rows***) before sampling (**A**), in leaf clips during Fv/Fm assay (**B, C**). Healthy and mildly stressed shoots (***green arrows***) when wet (***A, Low & High sites***), often appeared green (**A**) or blackish-red from protective pigments (***A, Low site***). Severe stress can appear orange-green (***A, High site***). Dead (chlorotic) shoots appear orange (***A, red arrows***). **B.** Subset of variously stressed shoots (chlorophyll fluorescence stress metric, ***Fv/Fm, listed for each sample***) from topography-shrub microsites (***labels***) by elevation zone site. **C.** Orange leaf tips suggest stressed shoots while Fv/Fm reveals healthy photosynthetic efficiency (Fig. 6), showing the importance of photosynthetic stress assays over visual stress surveys.

**Figure 6.**
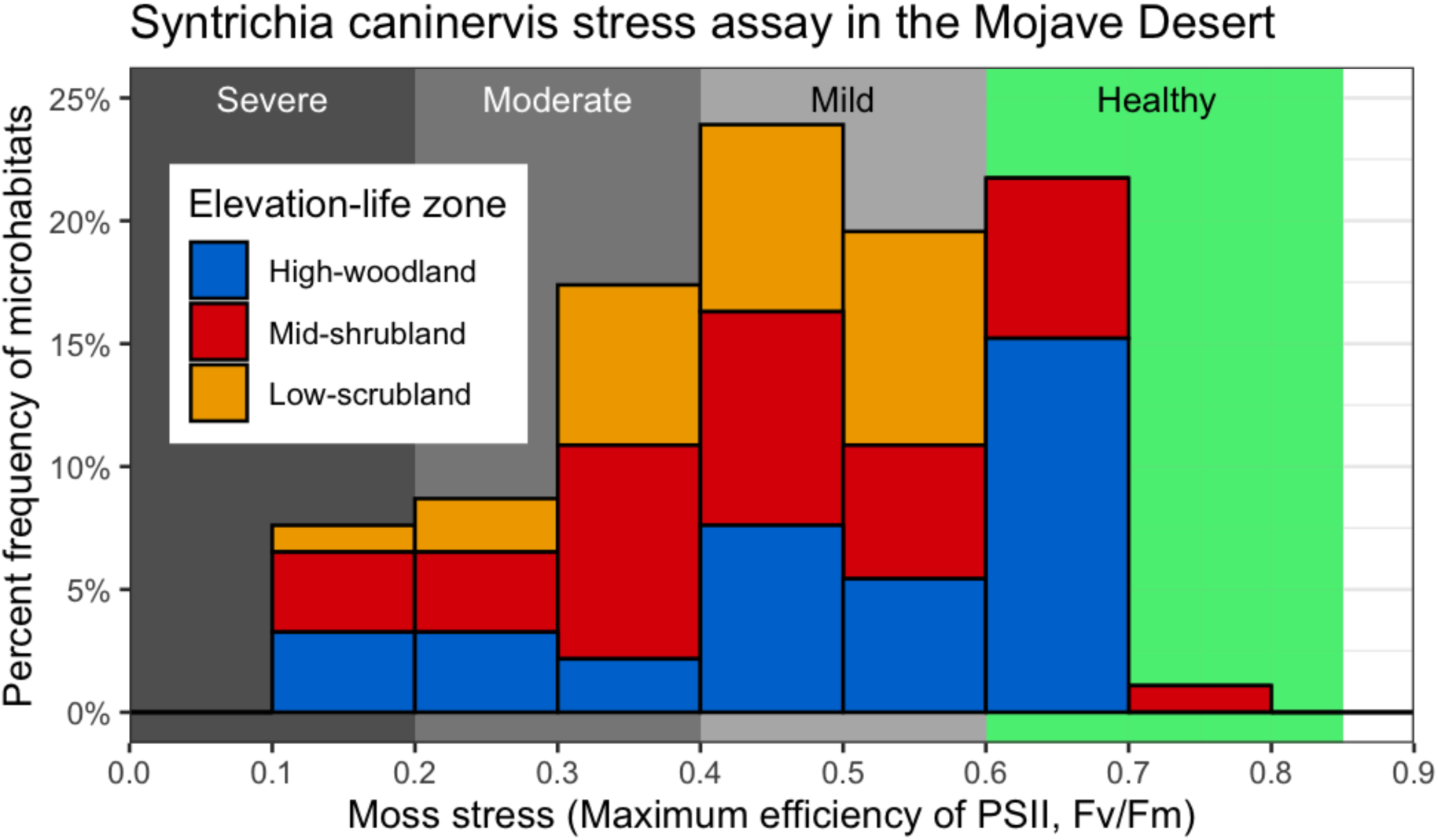
*Syntrichia* summer stress distribution by elevation zone site. Frequency histogram of biocrust moss (*Syntrichia caninervis*) microhabitats with severe, moderate, and mild levels of photosynthetic stress (estimated by the chlorophyll fluorescence metric, **Fv/Fm**) in 92 microsites colored by elevation zone site (see Fig. 1). Each bin is open on its maximum value (i.e. the bin [0, 0.1) does not include 0.1). Four colored bands indicate moss stress categories to aid climate vulnerability assessment in the Mojave Desert. Healthy mosses vary in their maximum Fv/Fm, so we have broadened “healthy” to begin at 0.6 (*see text*).

### Moss stress (Fv/Fm) vs three scales of habitat structure & buffering (Objective 4)

#### Habitat type ANOVAs

Contradicting our expectations, *S. caninervis* Fv/Fm did not differ on average by elevation zone (^Welch^F_2,59_ = 1.6, P = 0.204, **Fig. 4d**). Pooling topographical zones across sites, topography zone explained 21% of variation in Fv/Fm with the N-facing zone supporting healthier mosses, on average, than the S-facing or flat zones (^Welch^F_2,54_ = 12.7, P < 0.0001, **Fig. 4e**). Microhabitat type explained 23% of variation in Fv/Fm with healthier mosses under shrub canopies than in drip lines or interspaces (^Welch^F_2,45_ = 16.6, P < 0.0001, **Fig. 4f**).

#### Habitat buffering model

The full regression model including all multiscale buffering variables (elevation, PDIR, shade, and their interactions) was statistically significant explaining 55.4% of variance in Fv/Fm (F_7,84_ = 14.9, P < 0.0001; note this model should only be used for predictive purposes due to multicollinearity; **S2**). The highly significant reduced model (without elevation zone elevation) had acceptably stable beta estimates for PDIR, shade, and their interaction providing a model that can be reliably used for mechanistic inference (F_3,88_ = 23.5, P < 0.0001, R^2^ = 0.444, **Fig. 7**). Specifically, shade and PDIR were positively related to Fv/Fm (**Fig. 7b**); however, the full model indicated the strength of this relationship increased by elevation zone elevation (**Fig. 7a**).

**Figure 7.**
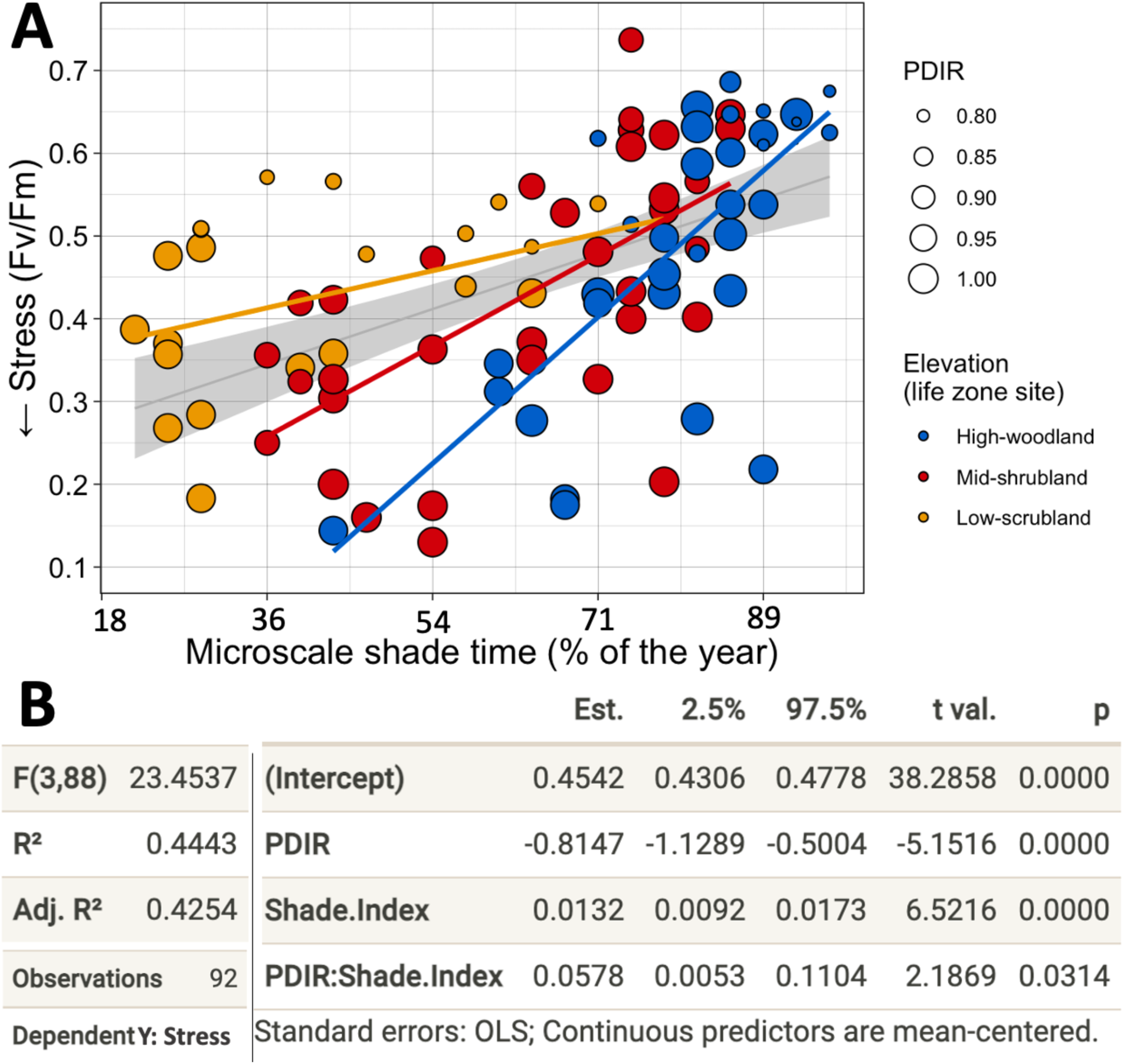
Summer stress vs habitat buffering proxies (elevation, PDIR, shade) **A.** Relationship of summer moss stress (lower Fv/Fm values indicate higher stress) of 92 *Syntrichia caninervis* microsites to habitat buffering proxies measured at three spatial scales: macroclimate (**elevation zone site: *point colors***), mesoscale potential direct incident radiation (***PDIR: circle size***), microscale shade time (***Shade.Index*** *in **B***), and their interaction (:). The OLS regression line for shade vs Stress is plotted with a 95% confidence interval in grey, and the interaction of elevation with shade is illustrated by varying slopes (*not tested*) of three OLS best fit lines plotted for each elevation zone. Elevation is shown graphically, but could not be included in the final model due to its multicollinearity with shade (see full model in **S2**). **B.** OLS regression results; residual plots are in **S3**.

#### Elevation zone models (topography zone and shade)

Fv/Fm was not related to shade at the Low-scrubland, but the two topography zones explained 57% of variation in which mean Fv/Fm was higher on the N-facing zone than on the Flat zone (^Wald^F_1,22_ = 26.9, P < 0.0001, **Fig. 8c**). At the Mid-shrubland, shade and topography and their interaction explained 67% of variation in Fv/Fm in which the Flat zone appears more positively related to shade than the N-and S-facing zones (*slopes not tested*; ^Wald^F_5,28_ = 12.5, P < 0.0001, **Fig. 8b**). At the High-woodland, only shade was positively related to Fv/Fm and explained 52% of variation (^Wald^F_1,32_ = 57.6, P < 0.0001; **Fig. 8a**).

**Figure 8.**
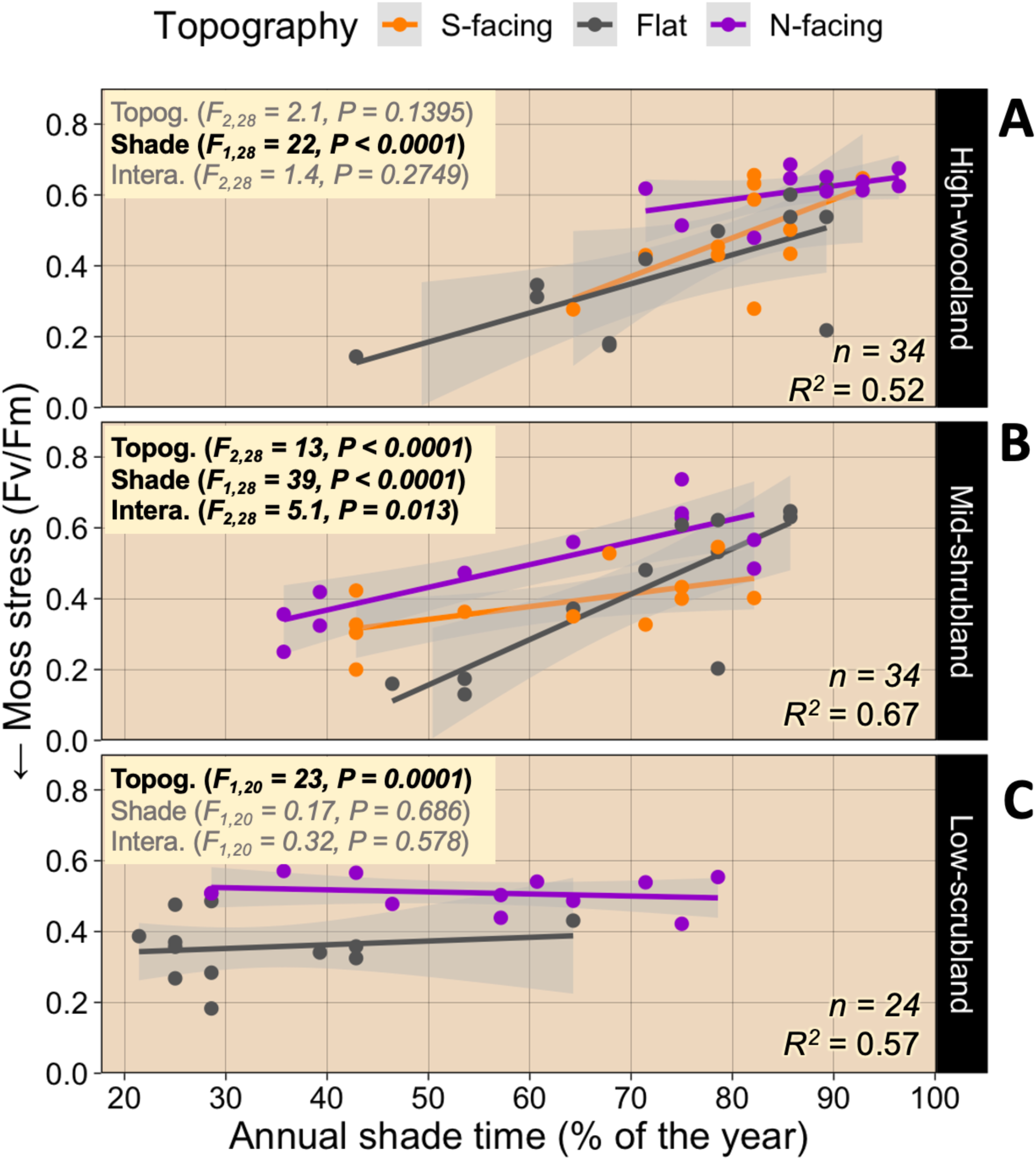
Summer stress vs topography and shade by elevation zone. Three elevation zone models and plots illustrate the linear relationship of summer moss stress of *Syntrichia caninervis* to topography (***Topog.***), annual shade time (***Shade***), and their interaction (***Intera*.**). HCCM-ANCOVA results for each elevation zone (***panels***) include alpha significance for each term in the full model (**bold *P < 0.05***), *R^2^* for the reduced model (including only significant terms), and best fit lines with 95% C.I. (***grey bands***) from OLS regression. ***Note***: no mosses occurred on the S-facing slope of the Low-scrubland (see Fig. 1c).

## DISCUSSION

### Fv/Fm summer stress interpretation

Our Mojave Desert stress assay confirmed the extreme resiliency of *S. caninervis* to present-day summer climate as we found no patch mortality in the 94 *S. caninervis* microsites studied (i.e. all Fv/Fm > 0, **Fig. 6**) despite the extreme weather of summer 2017 (**Table 1**, 2017), which included record-breaking temperatures (NOAA, 2020), low relative humidity, many small rain events, and over 45 days without rain. Moreover, only 33% of the *S. caninervis* microsites showed signs of moderate to severe photosynthetic stress as suggested by low Fv/Fm values relative to the maximum potential PSII efficiency of plants, ∼0.85 (**Fig. 6**). While over 50% of microsites had a moderate to extreme photosynthetic stress signal with severely stressed individuals present at all three elevation zones (**Fig. 6**). Fv/Fm can be confidently used as a stress proxy for desiccation tolerant mosses and revealed a much larger variation in photosynthetic summer stress and photoprotective acclimation than could be detected visually. The appearance of sampled patches ranged from obviously damaged shoots with reddish-orange leaf tips (severely stressed) to those lacking visible damage but darkly sun-pigmented from acclimatory zeaxanthins (Ekwealor et al. 2021; **Fig. 5**).

Reduced Fv/Fm in desiccation tolerant mosses may be caused by many stress factors including rapid drying, low water potential, high temperatures, excess light, and UV damage (Takács et al. 1999, Proctor 2001, Proctor et al. 2007, Greenwood et al. 2019, Ekwealor et al. 2021) all of which are likely acute or chronic stressors in these Mojave mosses especially during hydroperiods when metabolically active following rare precipitation events. Part of their impressive resiliency arises from being in the dry and dormant state most of the year during which they can tolerate extreme heat and extended drought (Stark 2017). When fully desiccated, *S. caninervis* can tolerate intense solar loading of 120°C (248°F) – two times greater than typical Mojave summer air temperatures (Stark 2005, Stark et al. 2009).

Historically called photoinhibition, reduced Fv/Fm has been described as a state of compromised efficiency in the light reactions of photosynthesis thought to result from deactivated PSII reaction centers and/or acclimatory changes in nonphotochemical quenching (NPQ; Demmig-Adams & Adams 2006). Specifically, there are three possible mechanisms of reduced Fv/Fm in desiccation tolerant bryophytes including rapid light acclimation through NPQ, and two longer-term acclimation responses including PSII repair cycling and slowly reversible NPQ (ql) which Muller et al. (2001) states are the result of prolonged, severe light stress. Regardless of the cellular mechanism, reduced Fv/Fm is usually interpreted as a photoprotective acclimation response to climate stress in mosses undergoing desiccation-rehydration cycles (Proctor & Smirnoff 2011, Proctor & Smirnoff 2000). Severely low Fv/Fm (< 0.2) can precede mortality (e.g. Coe et al. 2020), however, some experiments have shown recovery from such low Fv/Fm (e.g., Cruz de Carvalho 2011, Ekwealor et al. 2021). Further research is needed to determine in what climate scenarios does low Fv/Fm lead to reduced productivity or survival in bryophytes (e.g. Hájek & Vicherová 2013). Regardless of long-term resiliency potential, the large variation we observed in moss stress resistance presented a strong sample for testing the high-elevation and habitat buffering hypotheses.

### High-elevation refugia hypothesis

We had anticipated that the cooler, wetter climate and mountain topography at the High-woodland (**Table 1a**) would reduce summer moss stress relative to the Low-scrubland, but found no such evidence to support high-elevation buffering that could indicate potential for future high-elevation refugia in DNWR (**Fig. 4a**, **Fig. 6**). The lack of signal for high-elevation buffering along a 1200-m macroclimate gradient is surprising given the evident positive relationship between moss abundance and elevation along this aridity-stress gradient, the latter of which has been documented in other drylands for which floristics or community analyses have been conducted (Nash et al. 1977, Seppelt et al. 2016, Clark 2012).

In contrast, we found support for the habitat buffering hypothesis at all three elevation zones in which meso-and/or microscale habitat structure and buffering were positively related to summer stress resistance in *S. caninervis*. Mesoscale topography varied primarily by hydrological position and exposure (PDIR) while vegetation shading varied as a function of three microhabitat types related to shade shrub proximity (**Fig. 1c**). To our knowledge, this is the first study to link natural (i.e. nonexperimental) fine-scale habitat buffering to physiologically significant stress variation in a dryland moss. Future implications of these findings include the potential that the most resilient mosses may not simply be those at high elevations in drylands, but could be scattered throughout all elevation zones within species’ current ranges.

### Habitat buffering hypothesis

We next discuss how the fine-scale within-site habitat structure and buffering factors explored were related to moss stress in univariate and multivariate (i.e. multiscale buffering) frameworks.

#### Fine-scale habitat buffering

Supporting the habitat buffering hypothesis, high-shade canopy microhabitats were the most common habitat type for *S. caninervis* across elevation zones (**Appendix S1; see Supplemental Data with this article**) and harbored mosses with the highest Fv/Fm, thereby appearing the most stress resistant. Beneath these shrub canopies, mosses were shaded over 60% of the year regardless of elevation zone (**Fig. 4c**, **Fig. 4f**).

Furthermore, our novel shade buffering proxy, percent annual shade time (**Fig. 2**), appeared most important to summer stress resistance at the Mid-shrubland and High-woodland, where shade was positively related to Fv/Fm within each mesoscale topography zone (**Fig. 8a-b**).

However, we found no evidence for a shade-stress relationship at the Low-scrubland (**Fig. 8c**). The lack of a shade signal in this most arid and exposed elevation zone contradicted our expectation. Possibly, shade was overpowered by other habitat buffers; on the steep N-facing zone, topography buffering may overpower the buffering effect of shade, and hydrological buffering by the drainage basin may overpower shade buffering on the Flat zone where additional soil saturation may increase moss recovery time regardless of the shade level (**Fig. 1c**). Nonetheless, the patterns we present for fine-scale habitat buffering at the two higher elevation zones suggest that most shrub microhabitats in the Mojave Desert can provide an impressively high and physiologically meaningful shade buffer for *S. caninervis* regardless of shrub species and elevation zone.

In addition to microscale, mesoscale habitat buffering also appears important to moss stress resistance in DNWR (**Fig. 8**). We found small differences in topographical exposure (6-14% deviance in PDIR) are related to moss summer stress resistance and that, overall, northerly exposures appear to reduce stress even if slopes are gentle. For example, topography buffering appeared most important to stress at the Low-scrubland where mosses on the N-facing slope had significantly lower stress with exposure reduced by only 14% relative to the flat drainage basin (**Fig. 1c**, **Fig. 8c**). Mesoscale exposure may also be important at the High-woodland, but topographical exposure was confounded with shade here – the steep, N-facing zone was also the most shaded zone (**Fig. 8a**). Such ecological confoundedness is often unavoidable – greater shade on northerly-facing slopes is driven by topographical slope shading and higher vegetation density (Pelletier et al. 2017). However, we suggest that disentangling confounded fine-scale habitat features is less important for climate change vulnerability monitoring; determining the contribution of fine-scale vs. macroscale habitat structure to moss stress is perhaps a most critical first step to inform scales for monitoring and predicting future impacts on biocrust.

We found relationships that suggest mesoscale topography can drive unique buffering patterns for biocrust mosses that involve aspect and hydrological position that appear to influence biocrust moss distribution. For example, on the highly exposed S-facing slopes of the Low-scrubland. A mere 8% PDIR buffer (**Table 1c**) was associated with an absence of moss on our reference S-facing slope at the Low-scrubland. In fact, no mosses were found on any S-facing slopes in this elevation zone, suggesting a niche limitation for *S. caninervis* on soil surfaces receiving >1000 kJ/cm^2^/yr in low elevation Mojave Desert (**Table 1c**, **Fig. 1c**). Niche boundaries in response to such small differences in exposure shed light on the sensitivity of these poikilohydric organisms and corroborate prior warnings of their potential vulnerability to shifts in climate (Ladrón de Guevara and Maestre 2022).

#### Multiscale habitat buffering

We also found strong support for the multiscale habitat buffering hypothesis using multivariate models which revealed microsites for which two scales of habitat structure and buffering could predict moss stress resistance. At the Mid-shrubland, 67% of moss stress was predicted by mesoscale topography zone, microhabitat shade, and their interaction (**Fig. 8b**). Similarly, 43% of stress across all elevation zones was related to mesoscale PDIR, microscale shade, and their interaction (**Fig. 7**). There may also be an interaction between elevation zone and shade in which the importance of shade increases with elevation (**Fig. 7**). Overall, these models suggest that within-site fine-scale habitat buffering can have additive and interactive effects on moss stress at multiple scales and that potential exists for interactions with the macroscale, which could likely be in response to macroclimate (driven by elevation and shading by the dominant vegetation community). Specifically, it appears that effects of fine-scale shade are likely to vary based on topographical exposure, hydrological position, and elevation zone (or site) – thus we urge future researchers to anticipate such complexities in understanding moss climate stress resistance (**S2**).

In the Mojave Desert and other drylands, sheltered fine-scale habitats like N-facing slopes and soil beneath vegetation have been shown to reduce extremes in soil temperature, soil moisture, and moss thermal loading, but these studies do not link the habitat types or buffering to moss stress (Breshears et al. 1998, Bowker et al. 2000, Thompson et al. 2005, Kidron 2009). The only two studies that have linked natural variation to dryland moss photosynthetic stress also found microhabitat variation to be important, but did not compare its signal to macroscale environmental variation as we have done. The first study by Alpert (1988) used infrared gas analysis to compare carbon balance (i.e. the carbon offset between respiration and photosynthesis) of mosses cycling in and out of desiccation, and found healthier carbon budgets in N-facing, higher shaded habitats compared to S-facing, lower-shaded rock mosses in semi-arid California. The second study by Yin et al. (2017), experimentally altered shade environments in the temperate Gurbantunggut Desert of China. They measured healthier *S. caninervis* patches (e.g. greater Fv/Fm and antioxidant concentrations) under native shrub canopies compared to exposed patches after shrub-removal treatments. Our findings corroborate these studies and emphasize the potential that fine scale habitat buffering will be most relevant to these small plants if climate conditions persist or worsen.

### Can biocrust mosses hide from climate change?

Many studies predict that climate change will reduce dryland moss biomass and thus the functional roles of these plants in future drylands, which may cause ecosystem-wide consequences while potentially accelerating desertification (e.g., Coe & Sparks 2014, Rodriguez-Caballero et al. 2018, Ladrón de Guevara and Maestre 2022). Predicting such climate change responses in mosses is more complex than for other plants given their unique poikilohydric physiologies and fine-scale habitats, which require different temporal (e.g. intermittent hydroperiods) and spatial (i.e., microhabitat) scales of study than for large, homiohydric tracheophytes (He et al. 2016, Ladrón de Guevara and Maestre 2022). The shelter offered by shrubs and north-facing slopes in our study are examples of such fine-scale spatial complexity. These habitat features may have important implications for climate change presenting the likelihood that (1) microrefugia created by shade shrubs and northerly slopes will best protect biocrust mosses in future climates, and that (2) such fine-scale microrefugia will likely be more important to moss long-term resilience than macrorefugia existing within or outside species ranges.

Increased drought or erratic summer precipitation is predicted to increasingly stress sensitive moss hydration cycling (e.g. Coe et al. 2012). We warn that these hydrological stressors may strengthen the moss-nurse shrub dependency presenting an additional factor of vulnerability for biocrust mosses, as some shrub species may not be resistant to climate change. However, high-shade microhabitats in our study were not restricted to a particular shrub species by elevation zone, but rather, included over thirty species. This lack of shrub specificity should reduce future vulnerability in biocrust mosses due to the prevalence of diverse and abundant shrub habitat. We must warn, however, that *S. caninervis* establishment was most frequent under shorter shrubs with low-lying canopies (<0.5 m) such as *Ambrosia dumosa*, *Coleogyne ramosissima*, and *Artemisia* spp. at the Low-scrubland, Mid-shrubland, and High-woodland, respectively. Therefore, there is risk that large patches of biocrust moss may not survive if such low-lying dominant shrub species experience die-back (e.g., Ladrón de Guevara & Maestre 2022). Nonetheless, the previously mentioned niche shrub diversity may help alleviate impacts across many other shrub microhabitats.

Furthermore, unlike tracheophytes whose distributions are usually constrained to narrower elevational bands and single continents (Patiño & Vanderpoorten 2018), many moss species including in *S. caninervis* have broad ecological amplitude (BFNA 2007, eFloras 2023, María Ros et al. 1999), which is thought to be a function of their ability to exploit similarly buffered microhabitats existing across vastly different environments (Ladrón de Guevara & Maestre 2022) often cross-continental or disjunct locations (e.g. Brooks et al. 2023, Burge et al. 2016, Carter et al. 2016). The broad elevational and fine-scale niche we observed in *S. caninervis* in DNWR exemplifies such a pattern in which many shrub species appear to offer similarly buffered microhabitats distributed across at least three unique elevation zones.

Therefore, we foresee potential that the diversity of shade shrubs spanning multiple elevation zones and plant communities in the Mojave Desert may offer widespread and common microrefugia for biocrust mosses in future climates and may serve to minimize climate impacts on moss frequency, abundance, and ecological functions in biocrust.

## CONCLUSIONS

We used a natural experiment to test the habitat buffering and high-elevation refugia hypotheses for a broadly distributed, keystone biocrust moss, *Syntrichia caninervis*, within a climatically extreme part of its global distribution – the Mojave Desert. By characterizing its diverse microhabitat, aridity exposure, and photosynthetic stress resistance, we found evidence for physiologically significant summer habitat buffering in reduced-exposure microhabitats created by meso-and microscales of topography and vegetation sometimes interacting with each other or with macroscale elevation. These buffered microsites were distributed across all three elevation zones in the species’ current range, failing to support the high-elevation buffering hypothesis while strongly supporting the fine-scale (within-site) habitat buffering hypothesis. We conclude that many of these buffered microsites appear as candidate microrefugia potentially capable of “hiding” this species from future climatic extremes via sheltering by shrubs and N-facing slopes. Contrary to most climate change experiments predicting low moss resiliency to increasing aridity and altered precipitation patterns in drylands, our findings present the possibility that at least one ecologically critical biocrust moss may be more prepared for a changing climate than previously thought, as long as associated shrub mortality does not significantly accelerate.

## Acknowledgments

We thank the doctoral committee of TAC (Drs. Daniel Thompson, Sandra Catlins, Dale Devitt, and Peter Nelson) for assistance in the development of this University of Nevada Las Vegas (UNLV) dissertation project. We thank Amy Sprunger for logistical support in field research at the Desert National Wildlife Refuge. We thank Brian Bird for infrastructural support working within the NevCAN Network. Field sampling, lab work, and data management were aided by many volunteers to which we are grateful. The research was supported by the UNLV Summer Doctoral Research Fellowship and the American Bryological and Lichenological Society Anderson and Crum Field Research Award (to TAC).

## Author Contributions

TAC conceptualized this dissertation research under advisership by LRS and DD. TAC performed all analyses, made graphics, and wrote the original draft. TAC and AR performed field investigations. TAC, AR, and JG performed the laboratory investigation. All authors contributed to manuscript review and editing.

## Data Availability

Data and R code used in this manuscript are openly available on GitHub at https://github.com/TreesaClark/Syntrichia_summer_stress.

## Supporting Information

Additional supporting information may be found online in the Supporting Information section at the end of the article.

**S1.** Microhabitat type relative frequency by elevation zone site.

**S2.** Correlation and regression table for the full habitat buffering model.

**S3.** Residual diagnostic plots for the reduced habitat buffering regression model.

## APPENDIX

**Supplement 1.**
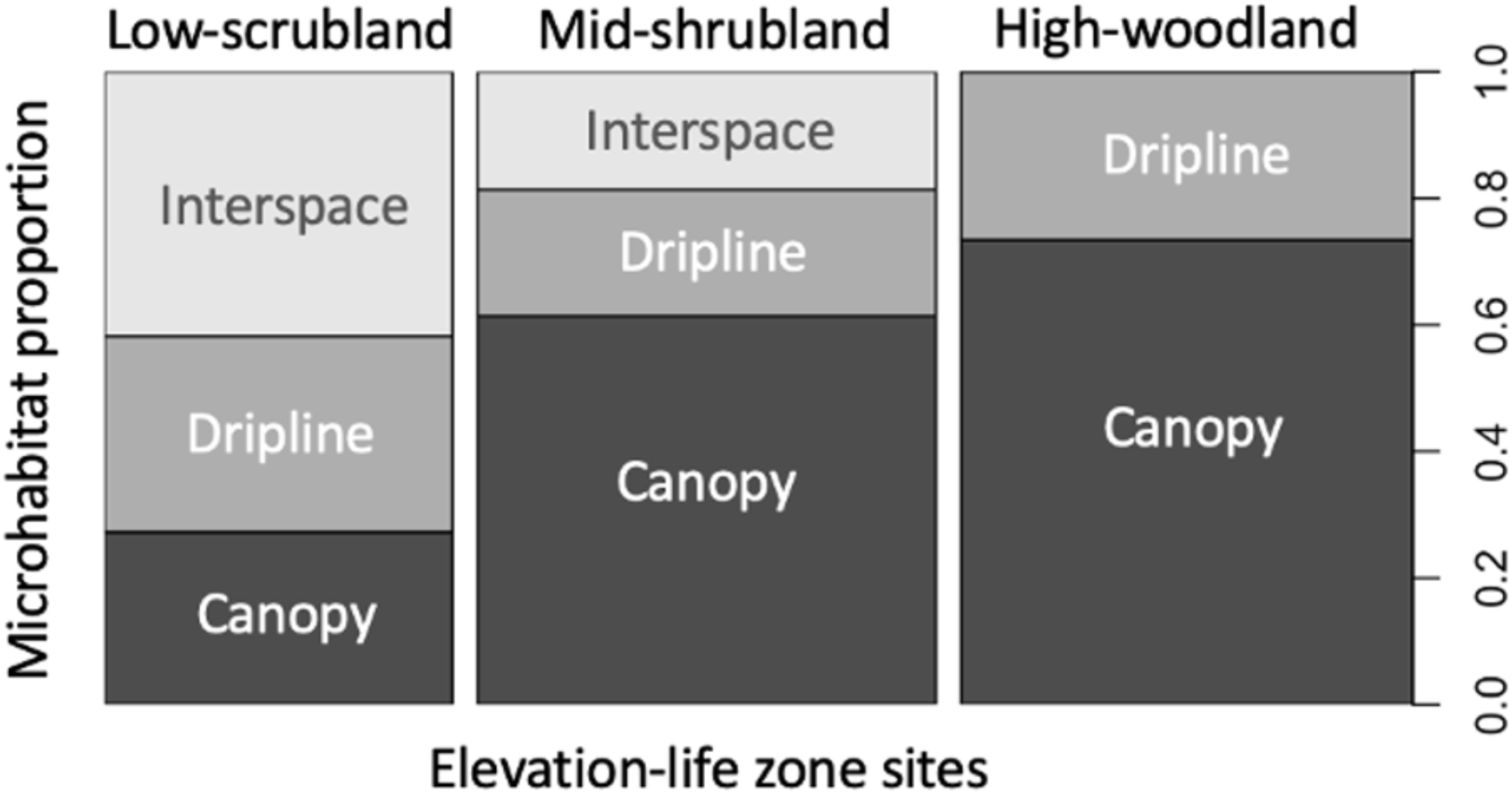
Microhabitat type relative frequency by elevation zone site. Relative frequency (proportion by elevation zone site) of three microhabitat types (shrub proximity classes: canopy, dripline, interspace) supporting *Syntrichia caninervis* in the DNWR elevation zone sites. Surveying for microsite distribution was conducted within a 2-km radius of each elevation zone climate tower and should be representative of elevation zone patterns at large in the Mojave Desert.

**Supplement 2.**
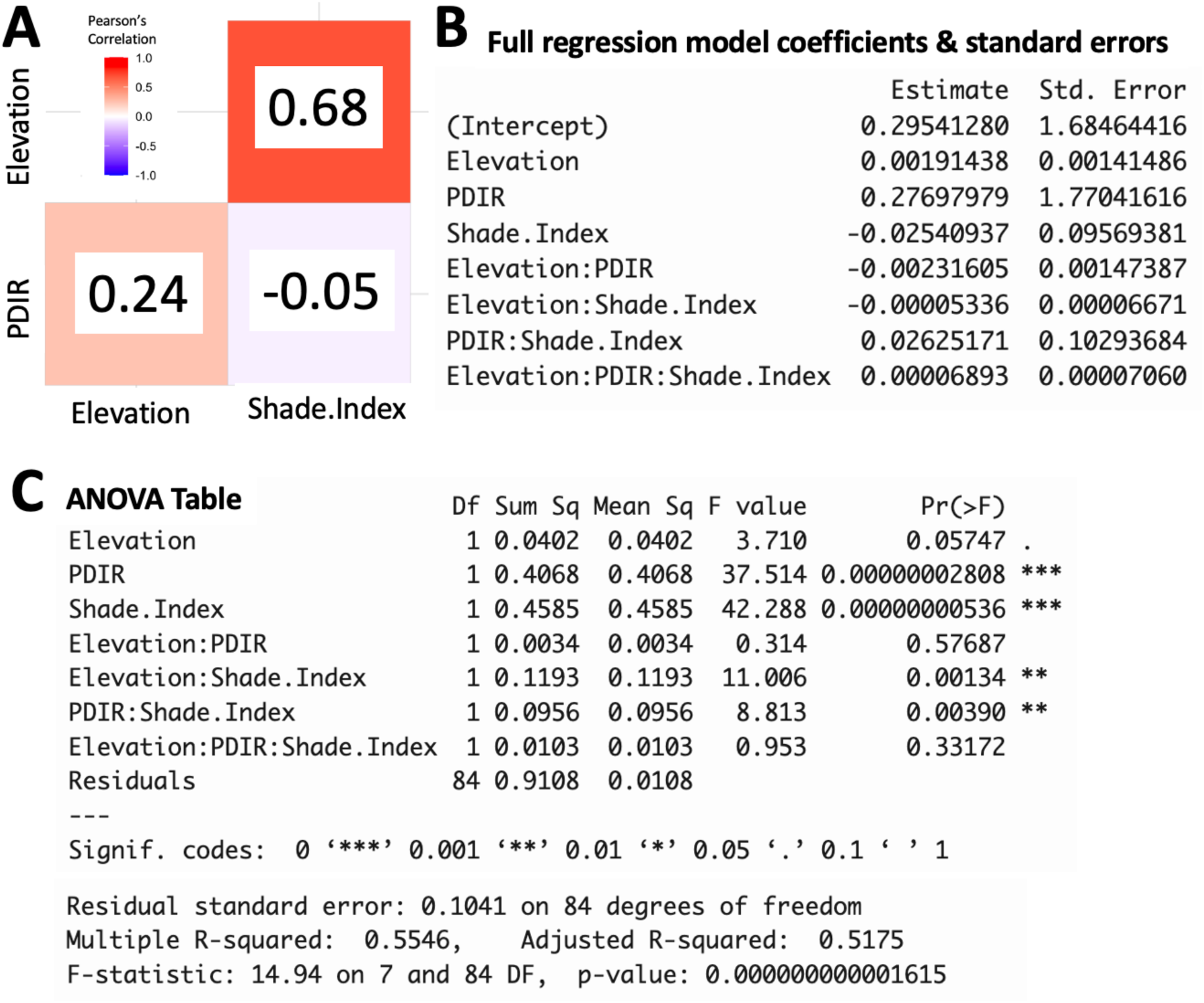
Correlation and regression table for full habitat buffering model. **A.** Pearson’s correlations table between regression predictors in the following models. **B**. Full OLS linear regression model of the relationship of summer moss stress (Fv/Fm) in *Syntrichia caninervis* and three habitat buffering proxies: elevation zone elevation (***Elevation***), potential direct incident radiation (***PDIR***), microscale shade time (***Shade.Index***), and their interactions. Elevation and shade suffer from multicollinearity such that this model is appropriate only for prediction rather than precise coefficient estimation (***see Analysis***). **C.** Corresponding ANOVA model (Type I errors) results, which do not suffer from multicollinearity; residual plots reasonably met assumptions and are not shown.

**Supplement 3.**
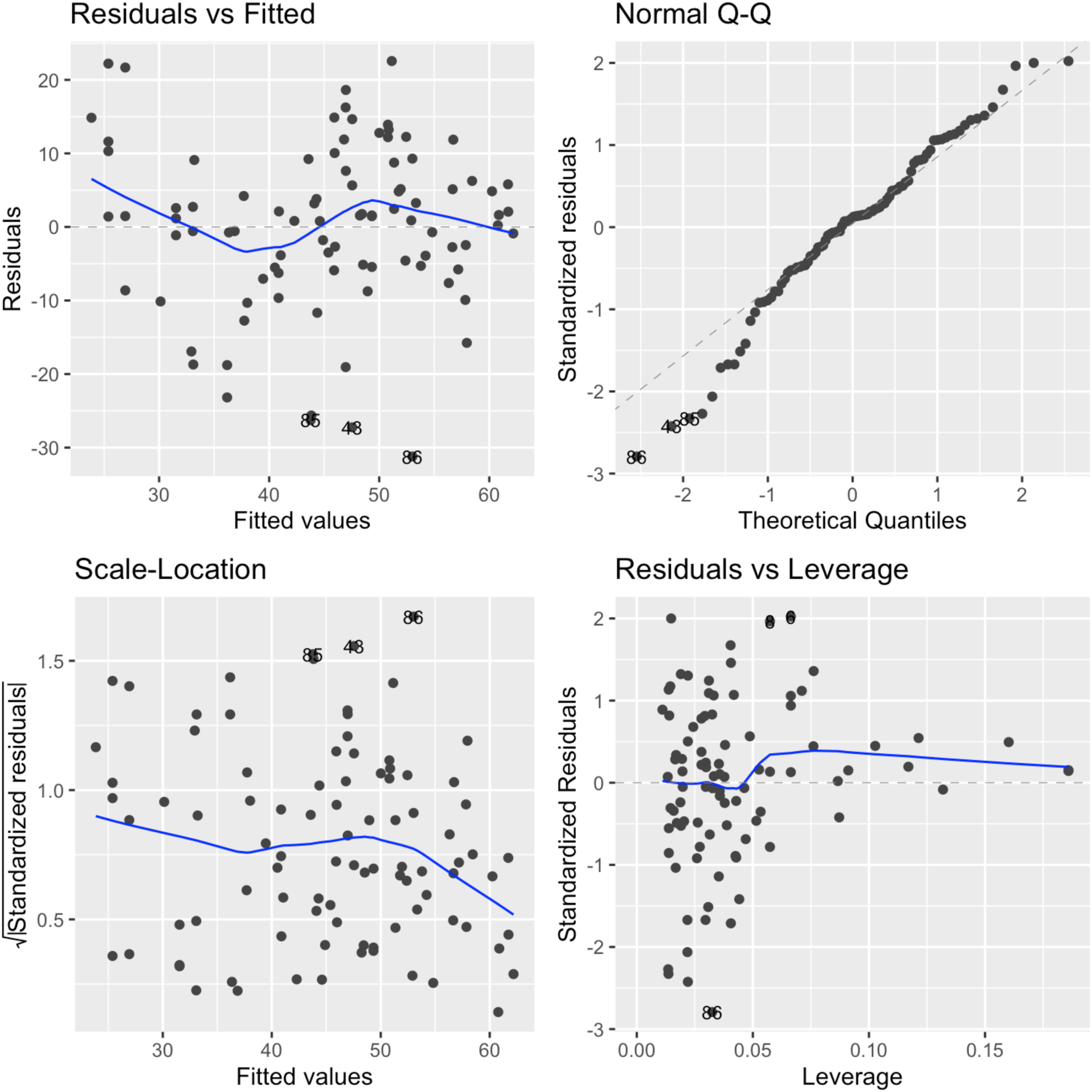
Residual diagnostic plots for reduced habitat buffering regression model. Residual diagnostic plots (*R stats::plot(Y∼X1*X2)*) for the OLS (ordinary least squares) regression model for moss stress (Fv/Fm) as a function of two habitat buffering predictors and their interaction term: potential direct incident radiation (PDIR) and annual shade time. See ***Analysis*** for details on model selection and **S2** for the full model. Model statistics and plot are in Figure 7.

